# Occult polyclonality of preclinical pancreatic cancer models drives in vitro evolution

**DOI:** 10.1101/2021.04.13.439717

**Authors:** Maria E. Monberg, Heather Geiger, Jaewon J. Lee, Roshan Sharma, Alexander Semaan, Vincent Bernard, Daniel B. Swartzlander, Bret M. Stephens, Ken Chen, Matthew HG Katz, Nicolas Robine, Paola A. Guerrero, Anirban Maitra

**Affiliations:** Department of Translational Molecular Pathology, The University of Texas MD Anderson Cancer Center, Houston, TX, USA; University of Texas MD Anderson Cancer Center Graduate School of Biomedical Sciences, Houston, TX, USA; Sheikh Ahmed Center for Pancreatic Cancer Research, The University of Texas MD Anderson Cancer Center, Houston, TX, USA; New York Genome Center, New York, NY, USA; Department of Bioinformatics and Computational Biology, The University of Texas MD Anderson Cancer Center, Houston, TX, USA; Department of Surgical Oncology, The University of Texas MD Anderson Cancer Center, Houston, TX, USA

## Abstract

Intratumoral heterogeneity (ITH) is a hallmark of cancer. The advent of single-cell technologies has helped uncover ITH in a high-throughput manner in different cancers across varied contexts. Here we apply single-cell sequencing technologies to reveal striking ITH in assumptively oligoclonal pancreatic ductal adenocarcinoma (PDAC) cell lines. Our findings reveal a high degree of both genomic and transcriptomic heterogeneity in established and globally utilized PDAC cell lines, custodial variation induced by growing apparently identical PDAC cell lines in different laboratories, and profound transcriptomic shifts in transitioning from 2D to 3D spheroid growth models. Our findings also call into question the validity of widely available immortalized, non-transformed pancreatic lines as contemporaneous “control” lines in experiments. Further, while patient-derived organoid (PDOs) are known to reflect the cognate *in vivo* biology of the parental tumor, we identify transcriptomic shifts during *ex vivo* passage that might hamper their predictive abilities over time. The impact of these findings on rigor and reproducibility of experimental data generated using established preclinical PDAC models between and across laboratories is uncertain, but a matter of concern.

## Introduction

Pancreatic Ductal Adenocarcinoma (PDAC) is a highly lethal malignancy, projected to be the second-leading cause of cancer-related death in the United States by 2030 [1, 2]. With a dismal 5-year survival rate of 10% [3] and few clinically meaningful therapeutic advances in recent years, the need for clinical and research advances is urgent. Preclinical models of PDAC represent essential prerequisites for advancing cancer research and experimental therapeutics in this lethal disease. The most commonly used and widely published preclinical models of PDAC that have been, and continue to be, leveraged in research laboratories globally are adherent cell lines distributed by the American Type Culture Collection (ATCC). For example, a perusal of current Google Scholar metrics demonstrates ~35,000 publications for PANC-1 and ~20,000 publications for MiaPaCa2, two prototypal ATCC cell lines nearly ubiquitous in every PDAC cancer research laboratory’s incubator. Included in many of these publications, and typically used as contemporaneous “control” cells in experiments, are two immortalized, non-transformed human pancreas-derived cell lines – so-called Human Pancreatic Ductal Epithelial (HPDE) and Human Pancreatic Nestin-expressing (HPNE) cells. While the reach of these preclinical models in furthering our knowledge of PDAC biology is extensive [2–14], several inherent assumptions have been made regarding the stability of the genomic and transcriptomic clonal architectures of these cells over time, including the fidelity to “normal” parameters for HPDE and HPNE during *ex vivo* passage.

The justifiably increased focus on rigor and reproducibility in scientific research studies has meant more stringent requirements for authentication of cell line resources, but the microsatellite assays typically used for confirmation of cell identity provide minimal insights into genomic and transcriptomic variabilities that might arise when the apparently identical line is propagated at different laboratories [15], or in disparate culture conditions, such as two-dimensional (2D) *versus* three-dimensional (3D) growth. A recent seminal publication by Ben-David et al [16] identified widespread genomic and transcriptomic variabilities in single cell clones isolated and expanded from the same parental cancer cells, which further translated into differences in drug responsiveness *in vitro*, suggesting a staggering level of pre-existing intra-culture heterogeneity within cancer cell lines. Although PDAC cells were not included in this analysis, it underscores the need for an in-depth benchmarking study using commonly used cancer and non-transformed “control” cells, in order to fully glean the extent of intra- and inter-culture heterogeneity that exists in these preclinical model systems.

In addition to the use of adherent cell lines, efforts at better recapitulating the biology of *in vivo* disease have led to the burgeoning adoption of patient-derived organoid (PDO) models as an *ex vivo* platform in the PDAC field. Encouragingly, early passages of PDAC PDOs maintain genomic and transcriptomic features of the tumor of origin, and harbor a gene signature of response to commonly used cytotoxic agents, enabling therapeutic prediction [17–20]. Nonetheless, whether the clonal architecture of PDOs remains stable over more prolonged *ex vivo* propagation, considering both the time in culture and the lack of extrinsic cues from the tumor microenvironment *in vivo*, remains less studied.

In this study, we perform an in-depth single cell genomic and transcriptomic assessment of clonal heterogeneity in a panel of established and globally utilized PDAC cell lines (Panc-1, MiaPaCa2, HPAF-II and BxPC-3), immortalized “control” cells (HPNE and HPDE), and in an independent PDO which is compared to its earlier passage prior to prolonged *ex vivo* propagation. We demonstrate that pancreatic cell lines – neoplastic and non-neoplastic - are comprised of remarkable sub-clonal heterogeneity at single cell resolution, which is unlikely to be detectable by conventional “bulk” profiling. Unexpectedly, HPDE cells harbor substantial genomic alterations and a transcriptome that essentially resembles cancer cells, questioning their use as a “control” line in research studies. We further demonstrate marked transcriptomic differences (including expression-based subtype classification) in microsatellite authenticated MiaPaca2 cells obtained from three independent sources. Another notable finding of our study is the observation of marked transcriptomic divergence (incorporating the appearance of distinct therapeutically actionable pathways) when adherent (2D) parental Panc1 cells are grown in 3D cultures. Finally, we describe the significant genomic and transcriptomic divergence of a later-passage PDO from an earlier passage, reiterating the importance of limiting the window for *ex vivo* therapeutic prediction and other biological experiments in this model type. Overall, our findings provide data-driven benchmarks for the limitations of the most commonly utilized preclinical models and platforms in PDAC research, with implications for the rigor and reproducibility of data generated in the *in vitro* setting.

## Results

### Single cell analysis identifies clonal heterogeneity of pancreatic cancer and non-transformed cells

We cultured six pancreatic cell lines, including three MiaPaca2 lines from independent sources and a pair of Panc1 lines subjected to differing growth conditions, for a total of nine samples analyzed with single-cell transcriptomic profiling (scRNAseq, 10x Genomics) and genomic copy number variant detection (scCNVseq, 10x Genomics). All samples were submitted for fingerprinting via MD Anderson Core Facilities prior to sequencing to confirm cell line identity. Visualization of the combined scRNA-seq data (77,068 cells) with UMAP [21] (**Figure 1A**) revealed substantial heterogeneity across the cell lines, including only partial overlap in apparently identical cells obtained from distinct sources or cultured under different conditions (*see later*). We first validated that cell lines of published epithelial or mesenchymal differentiation state maintained key marker identities [22] (**Figure 1B**). For example, we observed high expression of the epithelial transcripts *EPCAM* and *KRT8* in the previously described “epithelial” lines, HPAF-II and BxPC3, and conversely, the prototypal mesenchymal transcript *vimentin* (*VIM*) was essentially restricted to the previously described “mesenchymal” lines Panc1 and MiaPaca2. HPNE cells, which were originally derived from a *nestin* (*NES*) and *NOTCH2* expressing pancreatic progenitor cellular population [23], were confirmed to retain both markers **(Figure 1B**). Notably, HPNE cells, along with Panc1 and MiaPaca2, also demonstrated extensive *CD44* expression, underscoring the enrichment of putative stem-like cells in culture. We observed that cell lines that had undergone differing culture conditions (Panc1 2D, Panc1 3D) or were from distinct laboratories of origin (MiaPaca2-A, B, and C) were still more transcriptionally similar to each other than to the other cell lines (**Figure 1A**). To quantify transcriptional similarity, we computed phenotypic distance, defined as the distance between the samples in the diffusion space (*see Methods*). Upon projecting the distances onto a cladogram, we observed that all three culture variations of MiaPaCa2 cell lines fall into a common clade (MP2-A, MP2-B, MP2-C), while Panc1 cells, irrespective of growth as monolayer or in 3D, branch into their own clade (**Supplemental Figure 1)**.

**Figure 1:**
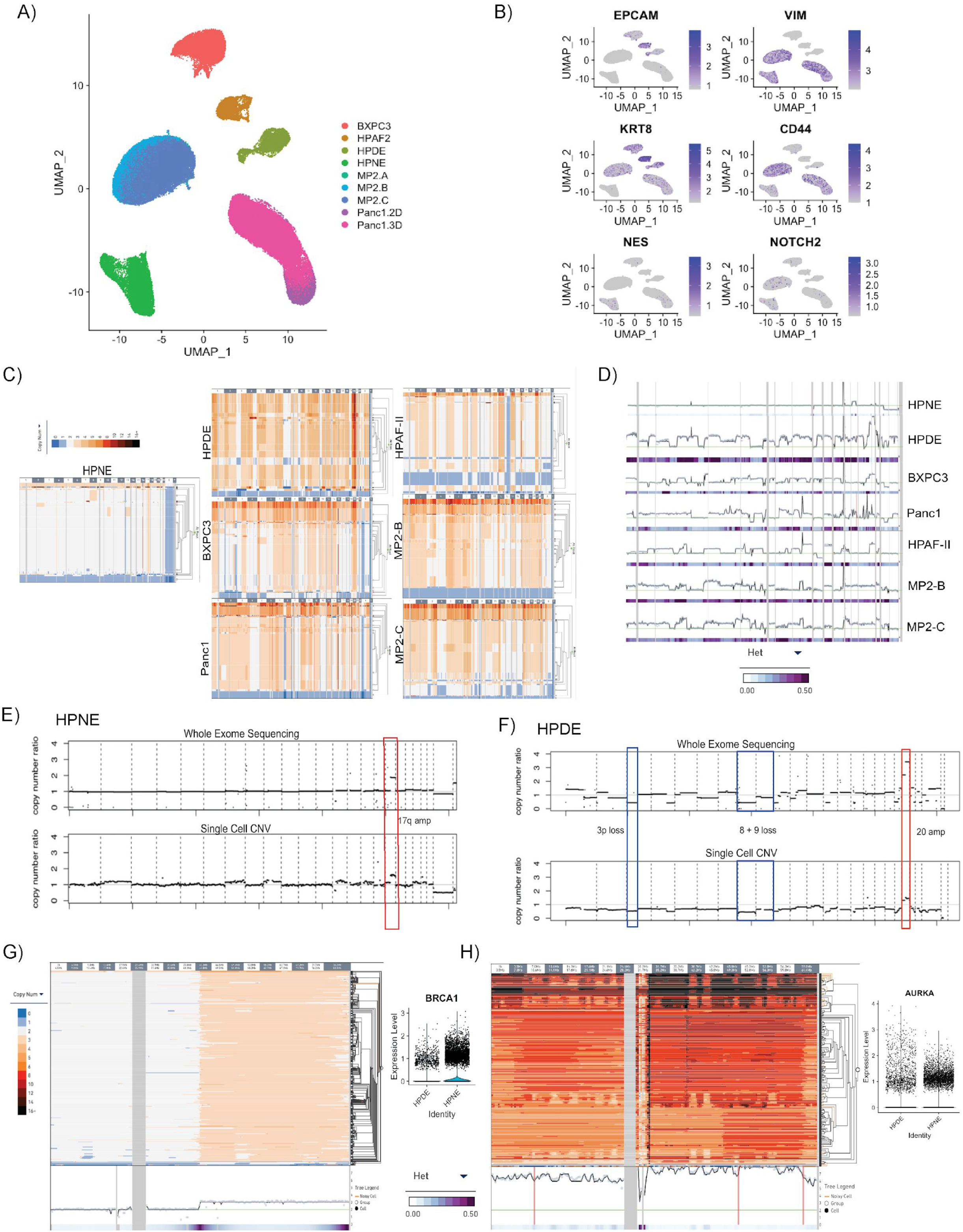
PDAC Cell Lines display heterogeneity at single-cell level. **A)** Uniform manifold approximation projection (UMAP) plot of single cells from PDAC cell lines used in this study. **B)** UMAP Feature Plotting for known cell-type markers (EPCAM, MUC1 for epithelial; vimentin, and KRT8 for mesenchymal cellular origin). **C)** Representative scCNV plots for cell lines. Columns indicate chromosomes, rows indicate individual cells. **D)** Segmentation plotting derived from ancestral node (max. nodes = 32) of scCNV clones from each cell line. Purple gradient indicates extent of deviation from ancestral profile across clones in a population. Columns indicate chromosomes, rows indicate ploidy. **E)** scCNV comparison to WES of HPNE cell line depicting amplified 17q as the only notable CNV event. **F)** scCNV comparison to WES of HPDE cell line. CNV events representing losses of chromosomal arms 3p, 8p, and 9p and amplifications at chromosome 20. **G)** scCNV high-resolution cell phylogeny of HPNE for all chromosome 17 locations showing ploidy = 3 for all cells at 17q arm. Corresponding scRNA data shows elevated expression of BRCA1, located within amplified HPNE region. **H)** scCNV high-resolution cell phylogeny of HPDE for all chromosome 20 locations showing ploidy >3 (as high as 13 in some cells at some locations) for all cells. Columns indicate chromosomes, rows indicate individual cells. Corresponding scRNA data shows elevated expression of AURKA, located within amplified HPDE region as a potential target of amplification.

We then performed single cell copy number analysis (scCNV) on the panel of cell lines using the 10x Genomics Cell Ranger DNA platform. The scCNV analysis demonstrated an overall striking degree of subclonal heterogeneity within each of the pancreatic cell lines, with dozens of clones bearing divergent genome-wide copy number alterations identified within each parental entity (**Figure 1C**). Using our “denoising” threshold of at least 20 cells/clone, we identified 19 distinct subclones in HPDE, 14 distinct subclones in Panc1, and 17 distinct subclones each in BxPC3 and HPAF-II (**Supplemental Figure 2**). Notably, of the two MiaPaca2 parental cells from distinct sources – MP-B and MP-C – that were assessed with scCNV, one harbored 18 discernible subclones and the other 20 subclones (**Supplemental Figure 2E and F**). Although 27 subclones were identified in HPNE cells, the majority of clones were highly similar to each other in terms of their scCNV profiles, and 30% of those cells sequenced segregate into designated Clone 4 (**Supplemental Figure 2A**), a fact reflected in the relative “tranquil” nature of HPNE whole exome and “pseudo-bulk” CNV data compared to HPDE cells (*see below*). We mapped the clonal families of each cell line to genes known to be amplified, deleted, or mutated in PDAC, and uncovered extensive heterogeneity in amplification and deletion profiles on a per-locus basis (**Supplemental Figure 2**). For example, the BxPC3 cell line, which is historically used as a *KRAS* wild type PDAC cell line, harbors copy number > 4 in nearly all clones at the *TERT* gene locus (**Supplemental Figure 2C**). Of interest, BxPC3 clone 2 (third largest clone detected, n > 200 cells) and clone 1 (n > 50 cells) harbored amplifications at the *BRAF* locus. It is known that in contrast to melanoma, cases of *KRAS* wild type PDAC, including the BxPC3 cell line itself, acquire in-frame deletions of *BRAF* that result in constitutive activation of the kinase domain [24]. However, amplification of the mutated “driver” oncogenic locus (e.g. *KRAS, BRAF, EGFR*) is not unusual, as it can provide additional survival and growth cues. Finally, the 10X scCNV data was converted into a “pseudo-bulk” copy number segmentation plot using the 10X analysis software (**Figure 1D**). The inferred ancestral clone was derived from the most parsimonious consensus explanation for each sample’s scCNV tree while keeping the number of allowable nodes constant. In the “pseudo-bulk” CNV data, previously described hallmark genomic events characteristic of PDAC lesions [1–6] were readily observed, albeit only in a subset of genomic subclones for each line. Specifically, amplifications of chromosomes 1q, 2, 3, 5, 7p, 8q, 11, 14q, and 17q, and losses of 1p, 3p, 6, 9p, 13q, 14q, 17p, 18q were distinguishable in segmentation plotting of the cell lines’ respective inferred ancestral clones, albeit showing a high degree of heterogeneity from the consensus profile at some loci (**Figure 1D**).

**Figure 2:**
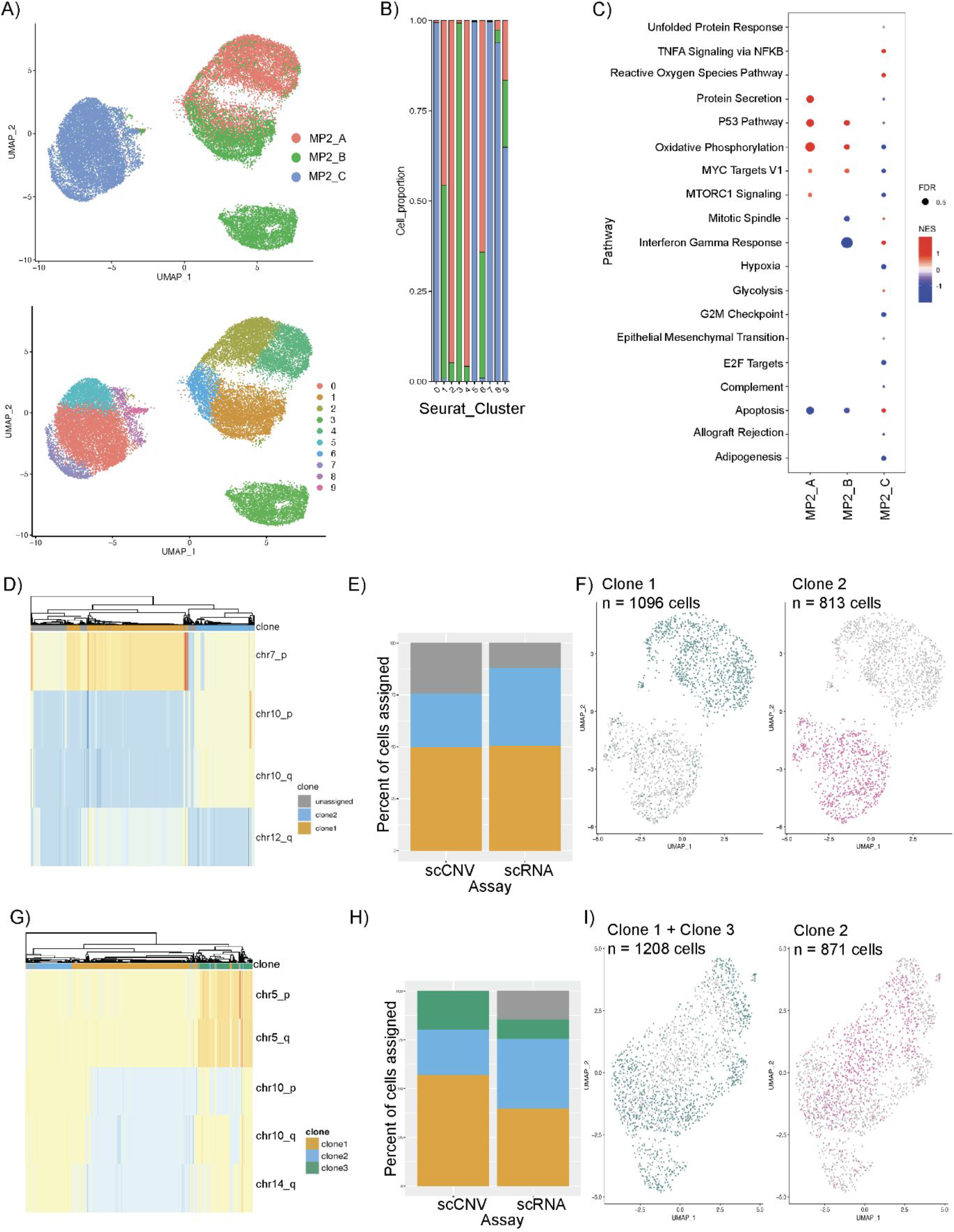
Characterization of custodial variability in MP2 cultures by scRNA and scCNV analysis. **A)** UMAP overlaying MP2 samples and their distinct clusters. **B)** Bar graph displaying distribution of cells per MP2 culture across clusters; MP2-A in red, MP2-B in green and MP2-C in blue. **C)** Bubble plot showing enrichment of pathways in MP2 cultures compared each other with normalized enrichment score (NES) on the x-axis. Size of the bubble represents false discovery rate (FDR); red indicates upregulation, blue indicates downregulation of the pathway. **D)** Genomic clones 1 (orange) and 2 (blue) identified from scCNV data of MP2-B based on CNV events at chromosomes 7, 10, and 12. **E)** Percentage of cells assigned to clones in scCNV and scRNA data. scCNV clones were mapped to corresponding scRNA data using CloneAlign. **F)** UMAP data of scRNA expression of single cells overlaid with CNV clones assignment for each cell; Clone 1 in green and Clone 2 in pink. **G-I)** Same as (D-F) for MP2-C.

### Single cell profiling reveals pitfalls of commonly utilized immortalized pancreatic controls

The Human Pancreatic Duct Epithelial (HPDE) cell line is a widely used model system for human pancreatic carcinogenesis [13, 25] typically as non-transformed, immortalized “control”. HPDE cells were derived from explanted human pancreatic ductal epithelium obtained from a pancreas resected for benign etiology, and immortalized using retrovirus-mediated expression of human papilloma virus 16 (HPV16) oncoproteins E6 and E7 [25]. The immortalizing effects of E6 and E7 are mediated primarily by targeting of the p53 and retinoblastoma (Rb) tumor suppressor genes [26]. Parental HPDE cells are wild type at the endogenous *KRAS* locus, non-transformed *in vitro*, and non-tumorigenic in mice, unless transformed through ectopic expression of mutant *KRAS*, in combination with other tumor promoting genomic events [27].

We thus hypothesized that the HPDE cell line would be transcriptomically distanced from the PDAC-derived lines, given its broad experimental application as a control line. However, our single-cell scRNA interrogation instead shows HPDE is most transcriptionally similar to HPAF-II, and on a distinct clade from the second commonly used “control” line, HPNE (**Supplemental Figure 1B**). On scCNV profiling, we find that HPDE cells retain a subclonal architecture akin to a neoplastic genotype in some clones (**Figures 1C and 1F**), while exhibiting a nearly genome-wide haploid state in the majority of cells sequenced. Conservatively, we identified 19 distinct genomic subclones in HPDE, with the largest subclones 12, 5, and 16 each representing more than 700 cells in culture (**Supplemental Figure 2B**). Segmentation profiles of the 19 clones demonstrated evidence of distinct amplification and deletions at loci linked to key PDAC-progressor genes. “Conventional” (bulk) whole exome sequencing (WES) on HPDE cells confirmed that it had undergone further bi-allelic losses of chromosomal arms 3p, 8p, and 9p, which are all “hot spots” housing known tumor suppressor genes in PDAC and other solid tumors (**Figure 1F, *top***), and this finding was validated in the “pseudo-bulk” CNV profile generated from scCNV data (**Figure 1F, *bottom***). Notably, all HPDE subclones harbor an amplification event (CN >10 in a minor fraction of subclones) at chromosome 20q (**Figure 1H**), bookending the *AURKA* gene locus within the most common region of overlap. Almost no heterogeneity was detected in the amplification event, reiterating that all HPDE single cells sequenced have undergone this amplification at some level (**Figure 1H, *purple bar***). Examination of *AURKA* transcript expression in scRNAseq data showed a spectrum of expression levels in HPDE, with a swath of cells expressing two-four fold higher expression of *AURKA* transcripts compared to HPNE cells (presumably representing the subclones with highest levels of genomic amplification). The encoded protein Aurora kinase A is commonly overexpressed in pancreatic cancers, and contributes to both chromosomal instability through destabilization of the mitotic spindle assembly and towards tumor progression via phosphorylation of substrate proteins [28, 29]. One can speculate that the widespread genomic perturbations observed in HPDE are at least partially a consequence of instability introduced by aberrant Aurora kinase A activity, in conjunction with the functional inactivation of p53 caused by HPV oncoprotein expression. To the best of our knowledge, this is the first description of *AURKA* amplification in HPDE, a purportedly “control” cell used in PDAC research, including in experimental therapeutics studies.

These findings in HPDE demarcate an important distinction from the second immortalized cell line commonly used as an experimental “control”, Human Pancreatic Nestin-expressing (HPNE) cells [23]. In contrast to HPDE, HPNE cells are non-ductal in derivation (as demonstrated by lack of epithelial markers, and widespread expression of progenitor cell transcripts *NES, NOTCH2* and *CD44* on scRNA Seq data, **Figure 1B**). In contrast to HPDE, HPNE cells were immortalized using retroviral transduction of the human telomerase reverse transcriptase (*hTERT*), which has been used in more recent times for derivatizing immortalized epithelial cells. Unlike HPDE cells, scCNV sequencing demonstrates that HPNE cells have not evolved a divergent subcellular taxonomy (heterogeneous deviation from ancestral clonal profile borders on a zero value, **Figures 1C and 1D**), with the exceptions of clones 9-12, 23, and 27, which comprise less than 15% of the total cell population (**Supplemental Figure 2A**). This scCNV finding is consistent with WES analysis of HPNE (**Figure 1E, *top***), which revealed a relatively neutral bulk segmentation profile, and is confirmed on the “pseudo-bulk” profile generated from scCNV data (**Figure 1E, *bottom***). While we do not observe amplifications or deletions of broad swaths of the genome, there is a discernible amplification event at chromosome 17q. As with HPDE, we interrogated genes contained within the amplified 17q locus, and *BRCA1*, among others, stood out as a candidate gene that also had corresponding scRNAseq overexpression **(Figure 1G**). The encoded Brca1 protein is a component of the homologous recombination repair (HRR) machinery, and *BRCA1* is uncommonly mutated in PDAC (~1%), typically in the germline, and associated with HRR defective cancers [30–33]. While the direct mechanism by which 17q would have been amplified remains to be investigated, we speculate that in the case of HPNE, the relative “genomic quiescence” reported here may be due, in part to the overexpression of *BRCA1* enabling DNA repair mechanisms. Ultimately, while HPNE does not exhibit the subclonal heterogeneity observed with HPDE, its lack of a ductal transcriptional profile and enrichment in markers associated with progenitor populations of uncertain histogenesis need to be factored in the use of these cells as an appropriate control for PDAC cells. To that end, the need for developing improved preclinical patient-derived “control” models for PDAC research is apparent from our findings.

### Custodial variability of MiaPaca2 cell lines drives transcriptomic heterogeneity

An identical parental line, when maintained under comparable passaging conditions in different laboratories (MiaPaca2), reflects a degree of custodial variability that impacts the translational value of that cell line as a controlled model. Previous work challenged the long-maintained assumption in cancer biology research that cell lines are clonal, stable entities, and explored the existence of phenotypic variability between widely used HeLa [15] and MCF7 [16] cell lines via cross-laboratory multi-omic comparisons. In such studies, cultured cells were maintained in uniform conditions after being obtained from different laboratory settings. Subsequent analysis revealed extensive inter-laboratory heterogeneity with respect to phenotype, copy number variations, and even drug response experiments, elucidating the high degree of variability underlying presumed “same” cell lines. Now, in light of single-cell analytics, we sought to investigate the extent of deviation from assumptions of clonality, both within and between cultures, in PDAC cell lines.

The MiaPaca2 cell line was initially generated from a primary PDAC lesion of a 65-year-old male, where the tumor had involved the pancreatic body, tail, and demonstrated periaortic invasion [11]. We assayed 3 “strains” of the MiaPaca2 cell line: one ordered directly from the American Type Culture Collection (ATCC), located in Gaithersburg, MD (MP2-A), a vial originating from an academic laboratory in Boston (MP2-B) and one that had been grown in our laboratory at MD Anderson Cancer Center (MP2-C). Following fingerprinting of the MiaPaca2 cells, we confirmed using ddPCR that all three versions of the cell line carried the same activating *KRAS*G12C mutation (**Supplemental Figures 3A-C**), as previously published in characterization studies of this cell line [11, 34, 35]. We report phenotypic divergence between these assumptively “identical” cultures that appears to be driven by changes at the RNA level. In our comparative scRNA-seq analysis of these samples, we find that the MiaPaca2 lines segregate mostly independently, with only a single small cluster where all three overlap (**Figures 2A, 2B**). MP2-A and MP2-B also share an additional region of overlap, but distinct from MP2-C. Gene Set Enrichment Analysis (GSEA) was performed to identify general pathways that are differentially altered between the samples (**Figure 2C**). MP2-A and MP2-B both share an enrichment in GSEA Hallmark pathways related to P53, oxidative phosphorylation, and MYC targets. Enriched in MP2-A are hallmark pathways related to mTOR signaling and protein secretion. In contrast, MP2-C has an almost entirely divergent GSEA profile, with enrichment in apoptosis pathways, interferon gamma, and TNF signaling.

**Figure 3:**
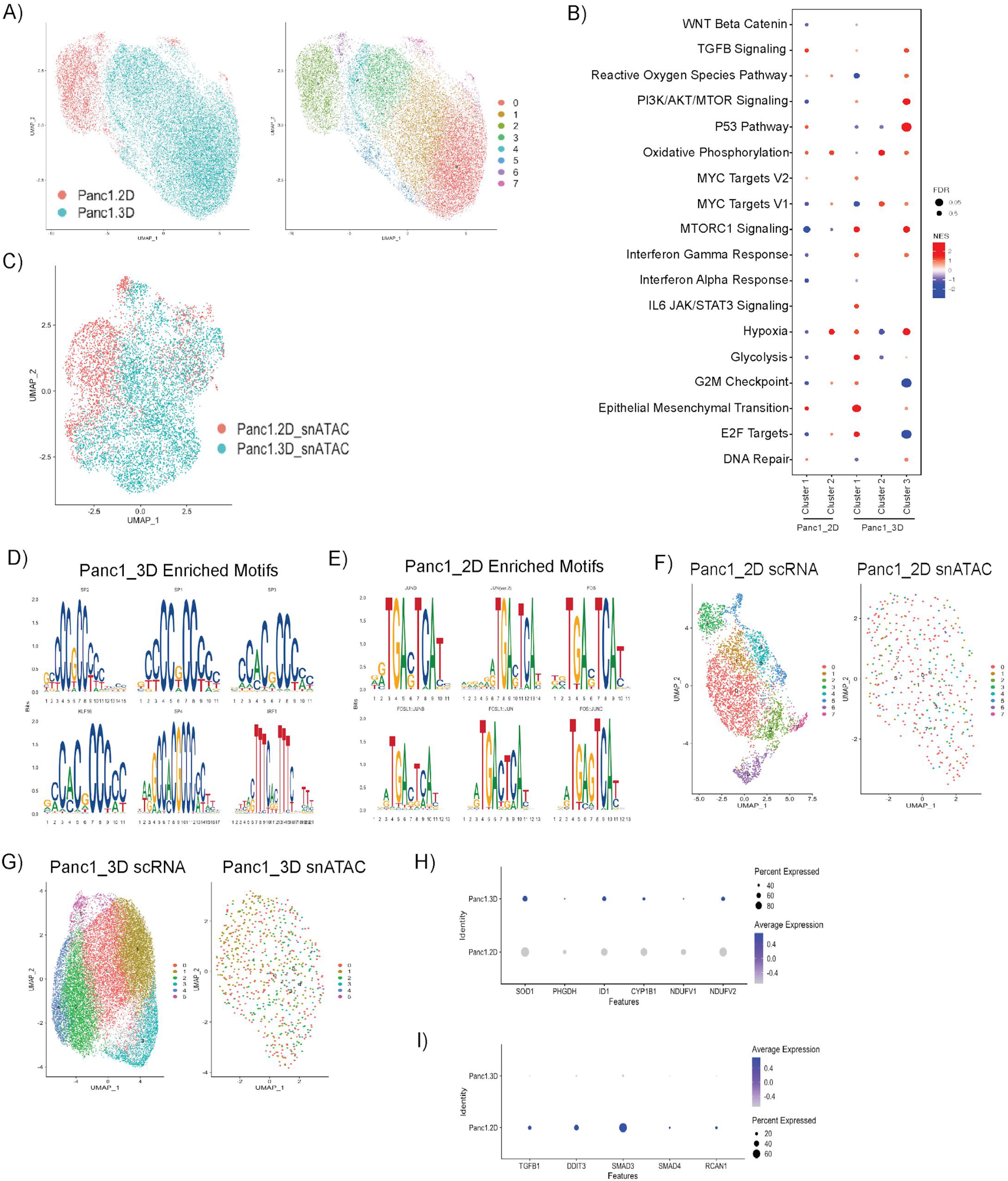
Spheroid growth model promotes transcriptional heterogeneity and epigenetic remodeling. **A)** UMAP overlaying scRNA of Panc1 samples and their unique clusters. **B)** Pathway enrichment profiles defined by GSEA analysis for Panc1-2D and Panc1-3D spheroid cell clusters. Size of dot represents FDR, red indicates upregulation, blue indicates downregulation of the pathway. **C)** UMAP representing snATAC-seq data of merged Panc1 samples. **D, E)** Analysis of merged Panc1 snATAC data shows enrichment in Sp/Klf motifs in spheroid data, and enrichment of Fos/Jun motifs in 2D monolayer culture. **F), G)** Integration of scRNA and snATAC data across both sample datasets. **H)** Comparison of expression of genes associated with Sp/Klf transcription factors in scRNA of Panc1 2D and Panc1 3D. **I)** Comparison of expression of genes associated with Fos/Jun transcription factors in scRNA of Panc1 2D and Panc1 3D.

Although we find evidence for a highly similar ancestral clone between the two strains (**Figure 1D**), on scCNV analysis, MP2-C has at least 2 additional detectable subclones (n = 20) compared to MP2-B (n = 18) (**Supplemental Figures 2E and 2F**). Further, these strains markedly differ in amplification and deletion events that correspond to chromosomal loci implicated in PDAC. For example, subclones in MP2-C have variable levels of amplifications at chromosomal loci containing the oncogenes *BCL6, BRAF*, and *PIK3CA1/2*, a pattern that is not observed in MP2-B. These initial findings demonstrate the remarkable heterogeneity, when viewed at the single cell level, across cell lines assumed to be derived from the identical parental origin.

### Evolution of divergent genomic subclones in MiaPaca2 strains has transcriptomic implications

While profiling at single-cell resolution makes evident the transcriptional and genomic heterogeneity of PDAC cell lines, integrating subclonal lineages and relationships between these independent datasets is crucial towards uncovering the impact of genomic alterations on transcriptomic diversity. Previous work applying statistical methodology and assumptions of copy-number gene-dosage effects on transcriptional expression [36, 37] have established a framework for how genome-transcription associations may be mapped in single-cell datasets derived from the same parental culture, but generated using individual assays [38, 39]. Using paired scRNA and scCNV from MiaPaca2 cells as a proof-of-concept for this method, we applied segmentation and copy number profiling from the scCNV data to clusters identified in scRNA transcriptomic profiles using the clonealign algorithm [40] (*see Methods*). An important caveat of this method is that inferCNV’s inherent design filtered out a large number of cells sequenced by scCNVseq. Thus, the findings discussed here are based on a subset of the more comprehensive subclonal families and profiles described earlier. Using clonealign filtering criteria to scCNV-derived MP2-B cells (**Figure 2D**), two distinct subclones were defined: cells harboring 7p amplification/10p/10q deletion, or cells lacking these CNV events. Of note, the “subsetted” clonal population lacking an amplification at 7p also had a deletion of the entire 12q arm. Applying the same clonealign filtering criteria to scCNV-derived cells to MP2-C (**Figure 2G**), three distinct subclones were defined: 1) with deletions of 10p, 10q, and 14q; 2) with no alterations in any of 5p, 5q, 10p, 10q, or 14q, and 3) with amplification of 5p and 5q, and deletion of 10p (but not 10q), respectively.

After the major subclonal populations were established from scCNV profiles, we used clonealign to identify subclones in the scRNA data that could be reliably mapped back to scCNV profiles (*see Methods*). Notably, we find that for MP2-B, the two unique genomic profiles we see in the scCNV data (**Figure 2E**) correspond almost perfectly to transcriptional subpopulations as visualized in our UMAP analysis (**Figure 2F**). Next, we used GSEA analysis to search for differences in expression signatures within each of the two delineated subclones. Subclone 1 of MP2-B (1096 cells) is characterized by four hallmark pathways, while MP2-B subclone 2 (813 cells) is defined by six, completely distinct gene sets (**Supplemental Excel File**). Given the difference in enrichment of cancer-related pathways that we found between the two MP2-B subclones, we then aligned MP2-B to MSigDB’s Oncogenic signatures, and found that subclone 1 is enriched for oncogenic pathways LEF1 upregulation, P53, and Cyclin D1, whereas subclone 2 is enriched for MTOR upregulation, EIF4E, RAF, NFE2L2, BCAT, and JAK2 (**Supplemental Excel File**). For MP2-C, we also see some degree of correspondence between the genomic profiles (**Figure 2H**) and the transcriptional subpopulations (**Figure 2I**), but the correlation is less striking than the one-to-one correspondence observed in MP2-B. In MP2-C, a very different pattern of enrichment is observed in both hallmark and oncogenic pathway sets. MP2-C subclones 1 and 3 were merged (due to very similar distribution across RNA transcriptomic clusters) for a total of 1208 cells, the population of which is significantly enriched for two hallmark pathways and five oncogenic pathways (**Supplemental Excel File**). Surprisingly, MP2-C subclone 2 (871 cells) is not significantly enriched for any hallmark or oncogenic gene sets (**Supplemental Excel File**). Importantly, there were also zero overlapping gene sets between MP2-B and MP-C, across any subclonal comparison. In summary, we confirm previous reports that progeny of the apparently identical parental cell line experience substantial genomic and transcriptomic divergence on prolonged *ex vivo* passaging, that such divergence is observed both within, and across, cultures of the parental cell line, and at least in some instances, the transcriptomic divergence can be ascribed to distinct subclonal genomic alterations.

### Modeling of tissue-based transcriptional PDAC subtypes using scRNA-seq data in established cell lines

Recently developed transcriptome-based subtyping of PDAC have demonstrated both predictive and prognostic significance in early studies. The dichotomous Moffitt classification schema of “basal-like” and “classical” subtypes was initially established using bulk RNA sequencing strategies applied to human tumor samples [18]. PDAC transcriptional subtypes are known to associate with specific microenvironmental niches [20], and primary tumors have been shown to undergo subtype switching in response to microenvironmental cues [41]. However, it is not well established whether cell lines can reliably classified into Moffitt subtypes [42] as freshly dissociated tumor tissue or PDO models, nor whether there would be an observable degree of subtype admixture when single-cell methods are applied to subclonal derivatives of parental cell lines. To investigate this, we sought to group the MiaPaca2 transcriptomic subclones into the dichotomous PDAC subtypes of the Moffit classification schema. We observed that MP2-A has all 9,768 cells represented by the Moffitt “basal-like” subtype, while MP2-B has subtype admixture, with 3,456 cells sorting as the “basal-like” subtype and 3,556 cells sorting as “classical”, and MP2-C has all 10,886 cells associated with the “basal-like” subtype (**Supplemental Table 1**). Upon further investigation, we found that these samples were being “forced” into one subtype or the other on the basis of a few genes being present or absent, as opposed to having a robust expression of the majority of genes specified by “basal-like” or “classical” subtypes. For example, MP2-A cells were sorted as “basal-like” based on the expression of *AREG* and *KRT15* (**Supplemental Figure 4A**), while MP2-C cells were sorted to “basal-like” based only on the expression of *AREG, GPR87, KRT15*, and *LEMD1* (**Supplemental Figure 4C**). Similarly, the MP2-B sample, for which we detected genes relevant for both “basal-like” and “classical” subtyping, we found that “classical-like” cells were classified as such based on only *LYZ*, while “basal-like” cells were classified as such due to expression of *AREG, GPR87, KRT15*, and *KRT7* (**Supplemental Figure 4B**). Given the thousands of cells present and the depth of sequencing on these samples, it is unlikely that this scant gene representation is due to dropout rates or data sparsity, which could result in skewed scRNA data. Rather, our data indicates that PDAC cell lines are sub-optimal models for application of tissue-based classification systems like Moffitt, largely because they might lack expression of substantial numbers of transcripts required for meaningful classification. This finding may impact how prevalent PDAC classification schemes are applied to monolayer culture models and any subtype-based therapeutic outcome predictions derived from these that are then extrapolated to patients.

**Figure 4:**
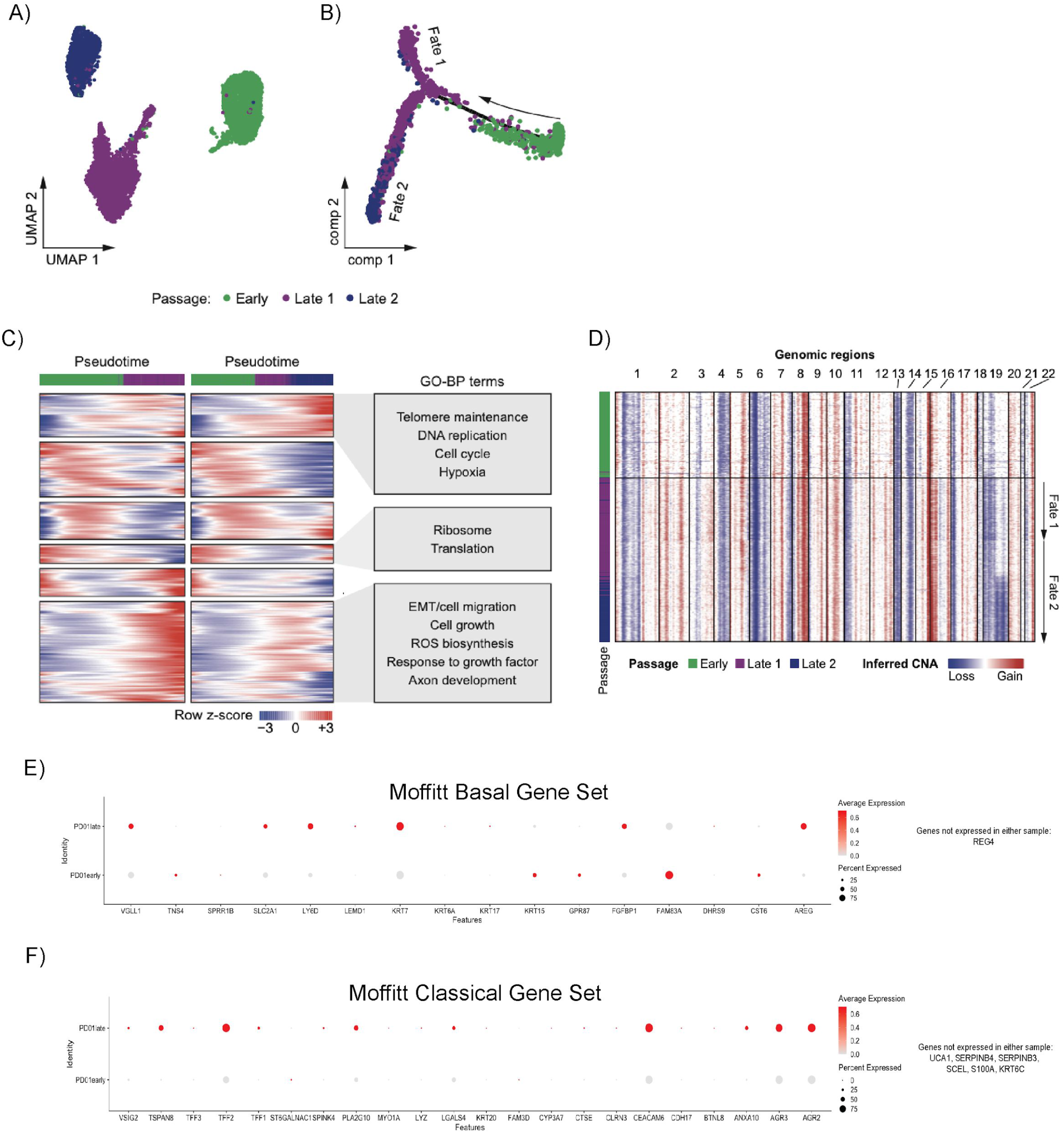
Patient-derived organoids evolve with time. **A)** UMAP plot of cells from a different PDO at early (green) and late (purple, navy) passages. **B)** Pseudotime analysis performed using Monocle of single cells from early (green) and late (purple, navy) passage organoids. **C)** Branched heatmap (left) showing dynamic gene expression along the pseudotime trajectory for each cell fate in B. Pseudotime progresses from left to right. Enriched Gene Ontology biological process terms for each gene cluster are listed on the right. **D)** Heatmap showing inferred copy number alterations of early and late passage organoids. Cells (rows) are ordered by pseudotime. Red represents copy number gains and blue represents copy number losses. **E)** Comparison of expression of Moffitt Basal subtype genes in scRNA of early (bottom) vs. late (top) PDO1. **F)** Comparison of expression of Moffitt Classical subtype genes in scRNA of early (bottom) vs. late (top) PDO1.

### Epigenetic alterations define transcriptional divergence observed in monolayer and 3D models

*Ex vivo* 3D models of tumor growth are being increasingly adopted, as a number of comparative studies between 3D spheroids and 2D monolayer cultures have shown that 3D culture more faithfully recapitulates *in vivo* biology [43–45]. Additional work has also shown that 2D monolayers might lack critical features associated with *in vivo* tumor progression, such as hypoxia [46–48]. Within the context of single-cell analytics, there is scant data to address the extent of transcriptional adaptations that occur in the process of modifying “conventional” monolayer culture to a 3D spheroid model. We find here that upon growth in 3D, Panc1 cells exhibit demonstrable transcriptomic divergence, as well as increased transcriptomic diversity, compared to matched 2D culture grown from the same parental vial, upon scRNA-seq analysis. While Panc1-2D and Panc1-3D cultures are overall more similar to each other compared to other cell lines at the transcriptome level (**Figure 1 and Supplemental Figure 1B**), they do not overlap when visualized using UMAP (**Figure 3A**). An unsupervised clustering shows that Panc1-3D cells organize into a larger number of clusters than Panc1-2D cells, highlighting greater intra-sample heterogeneity in Panc1-3D (**Figure 3A, *right***). To look further into pathway level differences between the 2D and 3D models, we computed differentially expressed genes and processed them with GSEA (*see Methods*). Panc1 spheroids upregulate transcripts related to PI3K/AKT/mTOR signaling and p53-related pathways, upregulate EMT-related transcripts in the largest subcellular cluster (**Figure 3B**: Panc1_3D Cluster 1), and downregulate glycolysis and E2F targets. Of note, differential regulation of PI3K/AKT/mTOR signaling in a 3D *versus* a 2D system (upregulated in spheroids, downregulated in monolayer) is consistent with prior reports [49].

As with MiaPaca2 samples, we sought to understand the basis of transcriptomic differences observed on scRNA-seq data, but using a slightly different analytic approach. As described earlier, spheroids were grown from the same parental population of Panc1 cells as the monolayer culture. scCNVseq was conducted on this parental population, and subsequent scRNseq + snATACseq was conducted after culturing the cells in their respective conditions. From the Panc1 scCNV data, we identified a total of 14 unique genomic subclones, albeit with some shared copy number aberrations amongst them (**Supplemental Figure 2G**). We then aimed to map this parental scCNV profile to the resulting scRNAseq data after growth under monolayer or spheroid conditions, to understand the subsequent transcriptomic reprogramming given identical parental genomic material. Applying the clonealign filtering criteria we used in our approach for MP2 cells, we defined two mutually exclusive subclones based on either 15q (subclone 1) or 14q deletion (subclone 2) for Panc1 (**Supplemental Figure 5A, 5B**). When we mapped these clonealign-defined unique genomic sublones to specific transcriptional programs across the two datasets, subclone 1 was comprised of 3,251 cells and subclone 2 of 404 cells in Panc1 2D scRNA-seq data (**Supplemental Figure 5C**), while in the Panc1-3D scRNA dataset, subclone 1 was represented by 904 cells, and subclone 2 by 223 cells (**Supplemental Figure 5D**). Remarkably, there was no overlap in the enriched hallmark or oncogenic gene sets for either of the two subclones between the two growth conditions, with only subclone 1 in Panc1 3D even demonstrating a substantial number of discernible gene sets (**Supplemental Excel File**). In other words, despite the common parental scCNV profiles, the transcriptomic divergence between the two growth conditions appears to be driven almost entirely by non-genomic events, such as epigenetic alterations.

Given the observational decoupling between Panc1 scCNV and scRNA datasets when comparing spheroids to 2D monolayer, we thus hypothesized that epigenetic modifications may be responsible for remodeling in the spheroid setting. To address this, we performed snATAC-seq and applied the Seurat analysis extension package Signac [50], across a population of 9,785 nuclei analyzed from Panc1 monolayer and 7,453 nuclei analyzed from Panc1 spheroids isolated from the same population, processed in parallel, as the cells used for scRNAseq. As a first measure of epigenetic heterogeneity, we find the spheroid nuclei in three different chromatin states (**Supplemental Figure 6A**) as opposed to 2 states observed in the Panc1 monolayer (**Supplemental Figure 6D**), indicating higher chromatin state diversity in the spheroid model. Further motif enrichment analysis within each population shows unique subcluster motif profiles (**Supplemental Figures 6B–6F**). We bioinformatically merged Panc1-2D with Panc1-3D nuclei to obtain a differential motif enrichment profile between the samples (**Figure 3C**). Notably, we observed enrichment of Sp/Klf-associated motifs in the spheroid model (**Figure 3D**), and Fos/Jun motifs enriched in the 2D snATAC-seq data (**Figure 3E**). We then mapped the snATAC-seq data to the respective scRNA-seq populations (**Figures 3F, 3G**), and confirmed the overexpression of putative target candidate genes affiliated with the enriched motifs that we had identified in corresponding snATAC-seq datasets (**Figures 3H, 3I**). In doing so, we find that the regulatory consequences of Sp/Klf motif enrichment are in genes associated with modulation of reactive oxygen species and metabolism-SOD1 [52], ID1 [53], NDUFV2 [54], and PHGDH [55]. The complete absence of expression, or, in some cases, downregulation, of these genes in the monolayer culture suggest that in the process of epigenetic reprogramming, Panc1 spheroids also undergo a full metabolic reprogramming by upregulating oxidative phosphorylation or glycolysis. This starkly different expression profile could have implications in the results of studies concerning metabolism in PDAC. In summary, our results suggest that the striking differences observed in scRNA-seq profiles between spheroid and monolayer growth systems are due to a divergence in the underlying epigenetic landscape, and the resulting differences in transcription factor activity and downstream pathways may have significant implications for the way these models are used interchangeably for *in vitro* experimentation.

### Long-term maintenance of organoids leads to transcriptomic change

PDOs have emerged as a promising preclinical model for therapeutic prediction and functional studies across multiple cancer types, including in PDAC [18, 19, 51–55]. Much like established cell lines, PDOs are maintained and expanded via *ex vivo* passaging in specially formulated “organoid” media. Early passages of PDOs retain the clonal genomic alterations observed in the parental tumor, as well as the transcriptomic signature(s) predictive of response to cytotoxic agents. However, whether the transcriptome of PDOs evolves over time in culture is less well studied. Therefore, we generated scRNA-seq profiles on a PDAC PDO at two timepoints, “early” and “late”, separated by 10 passages. We found that in the later passage of the PDO, two transcriptionally distinct clusters (**Figure 4A**) had emerged, which respectively formed two separate cell fates on trajectory inference (**Figure 4B**). Transcripts upregulated in cell fate 1 showed enrichment of epithelial-mesenchymal transition, cell growth, reactive oxygen species (ROS) biosynthesis, and axon development, while transcripts in cell fate 2 were enriched for telomere maintenance, DNA replication, cell cycle, and hypoxia pathways (**Figure 4C**). To test whether there were genomic alterations accompanying the transcriptomic divergence, we inferred copy number aberrations from the scRNA-seq data using the inferCNV package [56], which revealed a chromosome 16p gain in the later passage PDO, and an additional chromosome 19q loss specifically in cell fate 2 (**Figure 4D**).

Bearing in mind the emerging translational relevance of transcriptional subtyping of PDOs, we paneled for genes relevant to the Moffitt subtyping schema to understand subtype-specific shifts that may occur during *ex vivo* passaging. The earlier passage PDO has demonstrable expression of “basal-like” transcripts (**Figure 4E, *bottom***), but these did not overlap with the “basal-like” transcripts expressed by the later passage PDO (**Figure 4E, *top***). More striking was the differential expression of transcripts that define the “classical” subtype, which are expressed almost exclusively by the later passage PDO (**Figure 4F, *top***). We thus observe a subtype admixture over time, which is consistent with findings by Raghavan et al. [20], demonstrating that later passages of PDOs in culture undergo transcriptional reprogramming towards a “classical” phenotype, and reiterating the inherent plasticity of cellular subtypes. Of note, a recent assessment for “basal-like” and “classical” cells in PDAC tissues by subtype-specific protein markers has uncovered bi-phenotypic “hybrid” cells [57], while Ting and colleagues have shown evidence of subtype “drift” in PDAC tissues with various therapies [58], both lines of evidence supporting the plasticity of subtype states. As PDOs become increasingly adopted in translational research, the impact of transcriptomic reprogramming in later passages should be accounted for in the context of any predictive modeling studies.

## Discussion

As a globally utilized backbone of cancer research, established PDAC cell lines have played an indispensable role in enhancing our understanding of disease biology, and therefore, ensuring the accurate characterization of these tools is a prerequisite for ensuring downstream success in translational studies. Using in depth single cell analytics, we demonstrate striking subclonal heterogeneity in the most commonly used PDAC cell lines, identifying between 14 to 20 discernible subclones based on scCNV assessment. While established PDAC cell lines are assumptively oligoclonal based on published sequencing data in human tumors [59, 60], the level of subclonal heterogeneity observed on single cell analysis here is unexpected, leading us to propose the phrase “occult polyclonality” as the overarching genomic *sine qua non* of PDAC cell lines. Our data are comparable to the recent findings of Navin, Michor and colleagues, who identified between seven to 22 subclones on scCNV analysis of triple negative breast cancer tumors and cell lines [61]. Notably, these investigators also reported that cancer cells retain a “reservoir” of subclonal diversity, such that isolated single cell clones re-diversify their genomes within a relatively short time frame, at a calculated pace of one CNV event every four cell divisions. In line with this observation, we report that three independent strains of the same PDAC parental cell line (MiaPaCa2) demonstrably deviate from one another on single cell analysis, both in terms of underlying transcriptional profiles and the outgrowth of unique genomic subclones. As another concordant finding between the two studies, we confirm that “major” subclonal genomic events have a significant impact on transcriptomic divergence and clustering, not only within the altered chromosomal loci, but also more broadly in terms of enriched gene sets and pathways. The report by Navin, Michor and colleagues in breast cancer, and our own data in PDAC, harbors the potential to have a profound impact on the rigor and reproducibility of research, both within and across laboratories. For example, it calls into question the assumptions behind isogenic perturbation experiments in cultured cell lines, or the use of archived cell line specific profiling data for drawing inferences that are temporally disparate from the data generation point.

We further show that culturing a PDAC cell line typically grown in adherent monolayers as a spheroid model alters the underlying chromatin architecture and induces a more diverse transcriptional repertoire. In particular, we demonstrate acquisition of transcription factor programs in 3D culture that could confound the interrogation of therapeutic dependencies when compared to the identical cell line grown in 2D condition. While we do not posit that one culture condition is necessarily more reflective of *in vivo* biology than the other, it is noteworthy that certain dependencies (such as metabolic dependencies) initially identified in monolayer conditions have not subsequently been validated within an *in vivo* context. As model sophistication increases to more widespread adoption of the PDO format, we also report that, perhaps not unexpectedly, the molecular identities of PDOs tend to evolve over time, recapitulating what is observed with cell lines. In particular, our demonstration of selection against a basal phenotype in a PDO during *ex vivo* culture contribute to the growing body of evidence that therapeutic prediction studies germane to the parental tumor are best limited to early passages of derivative PDOs. While latter passages of PDOs can continue to serve as a *bona fide* preclinical model for PDAC research, they might need periodic re-interrogation with single cell analytics (or a reliable surrogate assay) to measure divergence over time.

Finally, we report here that the commonly used contemporaneous “controls” for PDAC cell lines, especially HPDE cells, are not actually as “normal” as current literature would lead one to believe. Previously undescribed in the literature, we identify considerable subclonal heterogeneity using single cell analysis (with an “occult polyclonality” that mirrors PDAC cell lines), and “major” chromosomal copy number events in HPDE and HPNE that translate to transcriptional upregulation of *AURKA* and *BRCA1*, respectively. Future studies may warrant a thorough investigation of whether omission of these chromosomal events in the context of comparative functional assays led to skewed results. Better yet, the systematic development, characterization, and dissemination of more reliable *in vitro* cell line “normal” controls for PDAC studies may be a requisite for maintaining the rigor of preclinical findings in the field.

## Acknowledgements

We thank L. Nakhleh and M. Edrisi for lending their bioinformatic expertise, and S. Muthuswamy for providing organoid growth media reagents.

## Methods

### Cell line selection and 2D culture

For non-neoplastic cell lines, HPDE and hTERT-HPNE were used, given their prevalence as “normal controls” in literature (over 800 citations for each line in PubMed). hTERT-HPNE was plated in Complete Growth Medium, consisting of 75% DMEM without glucose (Sigma D-5030) with an additional 2mM L-glutamine and 1.5 g/L sodium bicarbonate, 25% Medium M3 Base (Incell Corp. M300F-500), 5% fetal bovine serum, 10 ng/mL human recombinant EGF, 5.5 mM D-glucose (1g/L) and 750 ng/mL puromycin. HPDE cells were cultured using Keratinocyte Basal Medium with supplied supplements (Lonza, Clonetics KBM, Cat.CC-3111). For PDAC-derived cell lines, we wanted to ensure that lines selected were both widely-used in the field, and be representative of a range of characteristics in terms of differentiation and epithelial versus mesenchymal origin. MP2-A cell line was purchased from the American Type Culture Collection. MP2-B was obtained from a collaborating lab as a frozen stock, thawed, and plated upon arrival. MP2-C was thawed and plated from an in-house stock vial. All MiaPaCa-2 cells were cultured using DMEM (ATCC Cat. 30-2002) supplemented with 10% fetal bovine serum. BXPC3 and HPAF-II cell lines are commercially available, and cultured using RPMI 1640 (Gibco) supplemented with 2mM glutamine, 10% fetal bovine serum. Panc-1 cell line (commercially available) for 2D studies was cultured in DMEM with 2mM glutamine and 10% fetal bovine serum. Cell lines were harvested for single-cell dissociation and sequencing at 80-90% confluency. A volume of 3mL of media from each cell line was harvested and sent to the MD Anderson core facilities for mycoplasma testing. All cell lines were washed with PBS and incubated with 0.25% Trypsin for 3-5 minutes. For all lines except HPDE, trypsin was neutralized using each respective cell line’s media; in HPDE, trypsin was neutralized using a Trypsin Inhibitor (Invitrogen Cat. 17075-029). Cell lines were centrifuged for 5 minutes at 1200rpm, resuspended in cold PBS + 0.04% BSA (Roche Ref. 03116956001), filtered using Flowmi 40uM pipette filter tips (Sigma BAH136800040), and counted to ensure debris-free single cell suspensions. A portion of cells were sent to the MD Anderson core facility for molecular fingerprinting, and all cell lines were counted using a Countess II automated cell counter. 2,000 cells were loaded per lane into the 10x Chromium Platform for scCNV sequencing, and 10,000 cells per lane were loaded for scRNA sequencing.

### Panc-1 3D spheroid culture

For 3D spheroid growth, Panc1 cells from the same passage as the cultured 2D cells were thawed, washed with PBS, suspended in growth-factor reduced Matrigel (Corning #356231), and plated in 80uL spheres on 24-well Nunclon Delta surface plates (TMO140620). Spheroids were incubated for 10 minutes at 37° to allow the matrigel to harden, and 650uL of DMEM + 2mM glutamine and 10% fetal bovine serum were added. For spheroid dissociation in single-cells, media was removed, and spheroids were manually dissociated in 500uL of TryplE Express (Gibco Cat. 12604013). DMEM media was then added, and spheroids were centrifuged at 1500rpm for 5 minutes. To dilute the matrigel and obtain single-cells, cells were resuspended in 5mL PBS, filtered twice with 70uM cell strainers (Corning CLS 431751), resuspended once more in cold PBS + 0.04% BSA, filtered again using Flowmi 40uM pipette filter tips, counted, and loaded at 10,000 cells per lane into the 10x Chromium Platform for scRNA sequencing.

### Organoid generation

Dissociated cells were resuspended in PaTOM media (Huang 2015) and centrifuged at 1500rpm for 5 minutes. Cell pellet was gently resuspended in Matrigel Growth Factor Reduced (Corning) and plated as a dome in a Nunclon Delta Surface 24-well plate (Thermo Fisher). Cells embedded in matrigel were incubated at 37C for 15 minutes and covered with PaTOM media supplemented with Y-27632 dihydrochloride (Tocris)

### Single-cell RNA and CNV library construction and sequencing

All cell lines were processed using 10x Genomics platforms: Single Cell 3’ Reagent Kits v2 for scRNA and the Chromium Single Cell DNA Reagent Kit for scCNV. All libraries were assessed for quality on Agilent TapeStation 2200 and quantified using both Agilent TapeStation 2200 reagents and Qubit dsDNA HS Reagent kits (Invitrogen Cat. Q33231) before loading onto NextSeq500 (Illumina). Multiplexed scRNA libraries were sequenced using 150-cycle kits, paired-end, Read 1 26 cycles, Read 2 98 cycles, i7 index 8 cycles, i5 index 0 cycles. Multiplexed scCNV libraries were sequenced using 300-cycle kits, paired-end, Read 1 150 cycles, Read 2 150 cycles, i7 index 8 cycles, i5 index 0 cycles. Cell Ranger version 2.0 was used to convert Illumina base calls (bcls) to FASTQ files. FASTQ files for scCNV data and scRNA were aligned to the reference genome GrCh38 using files provided by 10x Genomics. Cell Ranger DNA Software v1.1 was used for subsequent downstream analysis of each cell line’s aligned data.

### Single-nuclei ATAC library construction and sequencing

Of the Panc1 monolayer and spheroid cultures, a subset of whole cells was isolated from the pools processed for scRNA + CNV sequencing. Using the nuclei isolation protocol and methods proscribed by 10x Genomics (Demonstrated Protocol CG000169, Rev D), we were able to load 5,000 nuclei per well for Panc1-3D and 7,000 nuclei per well for Panc1-2D for processing using the 10x Genomics Single Cell Atac Kit, v1. As with the previously described scRNA and scCNV libraries, all quantification and quality control was conducted on an Agilent 2200 TapeStation using D1000 High Sensitivity reagents, with additional quantification using Qubit dsDNA HS Reagent kits. A pooled library containing both Panc1-2D and Panc1-3D snATAC libraries was loaded on the NextSeq500 using 150-cycle kits, paired-end, dual-index mode, Read 1 50 cycles, Read 2 50 cycles, i5 index 16 cycles, i7 index 8 cycles. Cell Ranger version 3.1.0 and cellranger-atac version 1.2.0 were used to convert Illumina bcls to FASTQ files and align samples to reference genome GrCh38.

### scCNV Clonal Analysis

Using the Cell Ranger DNA software, maximum nodes were first set to 32 as per software guidelines to identify clonal tree roots and leaves (henceforth referred to as clones). Heatmap data for trees containing all leaves within the maximum node on a per-sample basis were exported (as an Excel file) and compiled so that cell counts/clone across chromosome regions could be quantified. A literature search was conducted to identify commonly amplified/deleted genes in PDAC, and the output of that search was combined with a Maitra Lab in-house panel of commonly mutated genes in pancreas cancers. Using the UCSF Genome Browser, chromosomal locations as provided by 10x Genomics software were mapped to gene-specific loci, and the resulting matrix of genes x clones x copy number was quantified. Clones containing less than 30 cells were excluded from the analysis to ensure residual sequencing artifacts would not skew the final result. Heatmaps were then generated using the heatmap.2 function in R studio.

### scRNA Seurat analysis of cell lines

Seurat (v3.1, Butler 2018) merged analysis was used to profile the scRNA cell line data. For removal of mitochondrial genes in cell lines, we subsetted all cells with less than 20% mitochondrial gene expression. Log normalization, variable feature identification (FindVariableFeatures), z-scoring (ScaleData) were applied to the merged object of all cell lines, and principal component analysis (RunPCA, npcs = 30) and subsequent dimension reduction (UMAP) were applied. Seurat’s FindMarkers command was used to identify cell lines that were representative of mesenchymal, epithelial, or neither phenotypes based on surface expression marker gene sets, as previously published by our laboratory (Castillo 2018).

### Gene Set Enrichment Analysis (GSEA) for MiaPaca2 Clusters

To obtain significantly enriched pathways on a per-cluster basis from MP2 cell lines, marker genes were extracted using FindMarkers function in Seurat. Resulting genes were ranked by −log10(FDR) multiplied by fold-change directions. This yielded an output wherein upregulated genes had positive values and downregulated genes had negative values. Pre-ranked GSEA was thus performed using fgsea R package v1.12.0 (Korotkevich 2019) specifying 10,000 permutations.

### snATAC analysis of Panc1 for motif activity, coverage plotting, and integration with matched scRNA

Using Seurat v3 and its extension package Signac version 1.0.0 (https://github.com/timoast/signac), along with additional R packages for gene annotation (EnsDb.Hsapiens.v86) and the JASPAR 2020 Motif Database (JASPAR2020), 10x genomic output files (metadata, fragments file, fragments index, filtered peak matrices) to generate Seurat objects containing motif information, gene annotations, and genomic ranges. Quality control on nuclei was done by quantifying nucleosome signal, TSS enrichment, number of peaks in fragmentation regions, and nucleosome signal according to Signac software recommendations. Samples were normalized using term frequency-inverse document frequency (TF-IDF) normalization, top features were identified, and dimensional reduction using singular value decomposition (SVD) on the TF-IDF matrix using peaks as features was done on both individual samples and a merged analysis object that had been created using fragmentation and peak data to identify common features to use as merging anchors. Motif activities were subsequently calculated using chromVar, as adapted for Signac. We then identified motifs that were enriched in each subcluster per sample, as well as differentially enriched between the two samples (merged analysis). To correlate regions of chromatin activity with scRNA data, per-sample snATAC-Seurat objects containing gene activity data were scaled, normalized, processed with latent semantic indexing (LSI), and UMAP dimensional reduction. We use all peaks that have at least 100 reads across all cells, and reduced dimensionality to 50, as recommended by Seurat guidelines. The processed snATAC dataset was then merged with the previously-described processing of scRNA data for Panc1 samples, with merging based on common anchors identified between the datasets, using scRNA data as a reference (FindTransferAnchors), to ensure commonalities and cluster structure between the datasets. Next, gene lists were curated using the GeneCards Human Gene Database-genes specifically activated by each enriched motif were queried and plotted using scRNA data (DotPlot).

### Cladogram analysis of cell lines

We sought to quantitatively assess the transcriptional similarity among the cell lines and establish if the families of origin for the cell lines are preserved. We use graph based methods called diffusion maps to compute the distances between the cell lines. In particular, we first map the cells from high dimensional transcriptomic space to a low dimensional rescaled diffusion space (Setty 2019), where phenotype distance between cells are more faithfully represented and compute distances between the cells in this new space.

Briefly, diffusion maps (Lafon 2006) is a family of techniques to reduce the dimensionality of any high-dimensional data and have gained a lot of application in the analysis of single-cell data (Azizi 2018, Mayer 2018, Haghverdi 2016). Essentially, they transform data from high dimensional gene expression space to a space of much lower dimensions, where usual Euclidean distance is reflective of the cellular phenotypic distance. Computing distances in diffusion space requires choosing the optimal number of diffusion coordinates. However, it has been proposed that rescaling the diffusion coordinates by their eigenvalues can circumvent this issue (Setty 2019).

For this, we combined all the samples and the resulting data was then reduced in dimensionality first using PCA. Diffusion Maps was then computed on the resulting PCA for which we first constructed a k-nearest neighbor (k = 30) graph using Euclidean distance and converted the resulting distance matrix into an affinity matrix using an adaptive Gaussian kernel (Setty 2019, van Dijk 2018). The resulting affinity matrix was row normalized (each row sum to 1) to obtain a Markov matrix, whose eigenvectors represent the diffusion coordinates. These coordinates were rescaled using the associated eigenvalues as described in (Setty 2019). Finally, we computed the Euclidean distance between every pair of cells between any two samples in the rescaled diffusion space. Thus, we defined the phenotypic distance between the two samples as the average distance between all pairs of cells as computed above. The resulting distance between samples was represented as a cladogram using the dendrogram function on linkage function in the scipy.cluster.hierarchy package in Python with parameter method = “average”. This was also represented as a heatmap using the clustermap function in seaborn package in Python.

### Phenotype map of existing gene sets

We sought to understand the intra-sample heterogeneity of MiaPaCa2 samples by doing subtype analysis. We utilized gene sets from Moffitt et al. (see main text reference) and used the expression of the genes to assign every cell to one of the subtypes. We constructed a simple assignment scheme to achieve this. We began by computing the average subtype expression for each single cell, i.e. average over the genes that define the subtype. We then reasoned that a single cell can be labeled as belonging to the subtype with the highest average expression.

### scCNV processing for scRNA clonal expression correlates

CNV calls per segment, per cell (from “node_cnv_calls.bed”) were intersected with mappable regions (as defined in “mappable_regions.bed”) using bedtools (v2.29.2). Next, the copy number for each mappable region for each cell was defined as the integer copy number with the most total bases covering the region. Finally, only regions of at least 2Mb on autosomes (chr1-22) were retained for downstream analysis. Cells defined as noisy in the QC summary file (“per_cell_summary_metrics.csv”) were removed prior to downstream analysis as well.

This data was converted to a cell x region matrix, with all integer values. Next, for each chromosome and chromosome arm, mean copy number was calculated by weighting based on the total length at each integer copy number, across all mappable regions. Ploidy was defined based on the median of these values across all whole chromosomes. Cells with ploidy more than one standard deviation from the mean across cells were removed.

Next, visual assessment of the chromosome arm mean copy number values per cell was used to determine which regions warranted further examination. Per sample, the chromosome arms that showed variation in a large percentage of cells were: MP2-B – chr7p, chr10p, chr10q, chr12q; MP2-C - chr5p, chr5q, chr10p, chr10q, chr14q; Panc1-2D – chr14q, chr15q. Next, all mappable regions in these chromosome arms were plotted for further visual assessment. In Panc1-2D, chr15q mappable regions starting at coordinates <40Mb or >80Mb, which showed a different profile from the other mappable regions, were also excluded. In MP2-B, only mappable regions from 93.5-126.3Mb on chr12q were considered, as the remaining regions had consistent copy number across cells. As all examined cell lines appeared to be triploid overall, deletion defined as mean copy number < 2.75, while amplification defined as mean copy number >= 3.75. Next, clones were defined as described in the main text. The end result of the scCNV workflow for each sample was a segment x clone matrix, where the segments were the mappable region coordinates and the values were integer copy number values.

### scCNV-scRNA integration

The mappable region coordinates were intersected with the gene coordinates (from Ensembl v84, same as in the CellRanger reference) to produce a gene x clone matrix. This matrix was used as input to clonealign (v2.0), which was used to assign cells in the RNA to each clone. Default parameters (including min total nUMI per gene of 20 and min total nUMI per cell for all copy-variant genes of 100) were used to filter genes and cells before running clonealign. Next, clonealign was also run with default parameters, except for the minimum probability required to call a cell as assigned to each clone. This parameter was set to 90% for MP2-B and Panc1-2D. For MP2-C this was lowered to 50%, due to very low numbers of cells with high confidence clone calls found in this sample.

Seurat (v3.1) was used for dimension reduction (UMAP). For this analysis, all annotated genes expressed in at least 10 cells were included. Next, only cells assigned to a clone with confidence above the threshold were included for downstream analysis. Finally, standard analysis was performed including SCTransform for normalization and variable gene selection, followed by principal component calculation on the variable genes (RunPCA). The first 30 PCs were used for input into UMAP (RunUMAP) for the MiaPaca2 samples, while the first 50 were used for Panc1.

Seurat’s FindMarkers command was used for differential expression between clones. For MP2-C, clones 1 and 3 were combined into one group to create a pairwise comparison versus clone2. This was because cells called as either of these clones showed very similar expression. All default parameters were used except for reducing the log-fold-change threshold to 0.10 from 0.25.

### GSEA of clonealign-defined scRNA clones

Pre-ranked GSEA was thus performed via the GSEA software version GSEA_4.0.3. Molecular Signature Databases h.all.v7.1.symbols_1.gmt (hallmark pathways) and c6.all.v7.2.symbols.gmt (oncogenic signature gene sets) were both used to align gene lists extracted from scRNA clones in Seurat, as previously described. A false discovery rate (FDR) cutoff of 25% was used to analyze the results for significant pathway enrichment per sample.

### Dimensionality reduction and cell clustering of PDO Pair

Initial single-cell analysis was performed using Seurat v3.1.0 (1). Cells containing less than 200 genes were removed, and log-normalized expression of 2,000 variable genes were used to reduce the data into two-dimensional space. Cells from PDO1 early and PDO1 late samples were combined using FindIntegrationAnchors and IntegrateData functions. To retain the highest number of cells for subsequent analyses, no additional filters were initially applied. Cells with higher expression of mitochondrially encoded genes were filtered out *post hoc* based on the results of the clustering.

### Cell state trajectory analysis of PDO Pair

Trajectory inference was performed using Monocle v2.13.0 (2). Analysis was done using raw counts obtained from Seurat objects with preserved cellular metadata. Top 1,000 variable genes were used to reduce the data and order the cells. Pseudotime heatmap was created using genes with dynamic expression changes obtained using differentialGeneTest function specifying fullModelFormulaStr = “~sm.ns(Pseudotime)” and q-value cutoff of 1 x 10^−40^. Branched pseudotime heatmap was created using BEAM function and q-value cutoff of 1 x 10^−5^. Genes were clustered based on pattern of expression changes and extracted from Monocle object for subsequent analysis.

### Copy number inference from scRNA-seq

Copy number alterations from scRNA-seq were inferred using inferCNV R package v1.0.4 (https://github.com/broadinstitute/inferCNV). Raw gene counts from filtered cells were used, and cutoff was set to 0.1. Stromal cells were used as reference.

## MAIN

**Supplementary Table 1.** Distribution of PDAC scRNA cells across Collisson + Moffit PDAC subtypes. Cell Count / Percentage of total.

| Cell Line | Moffitt- Basal | Moffitt- Classical | Collisson- Classical | Collisson- Exocrine | <u>Collisson- Quasimesenchymal</u> |
| --- | --- | --- | --- | --- | --- |
| MP2-A | 9768 / 100% | 0 / 0% | 4348 / 44.51% | 0 / 0% | 5420 / 55.49% |
| MP2-B | 3456 / 49.29% | 3556 / 50.71% | 2408 / 34.34% | 1896 / 27.04% | 2708 / 38.62% |
| MP2-C | 10886 / 100% | 0 / 0% | 3276 / 30.09% | 3724 / 34.21% | 3886 / 35.70% |
| BXPC3 | 5606 / 54.88% | 4609 / 45.12% | 3532 / 34.58% | 2175 / 21.29% | 4508 / 44.13% |
| HPAF-II | 2367 / 51.27% | 2250 / 48.73% | 1777 / 38.49% | 961 / 20.81% | 1879 / 40.70% |
| HPDE | 2879 / 55.41% | 2317 / 44.59% | 1594 / 30.68% | 1479 / 2.46% | 2123 / 40.86% |
| HPNE | 6318 / 49.80% | 6370 / 50.20% | 518 / 4.08% | 5221 / 41.15% | 6949 / 54.77% |
| Panc1-3D | 7440 / 51.03% | 7139 / 48.97% | 4883 / 33.49% | 3537 / 24.26% | 6159 / 42.25% |
| Panc1-2D | 2891 / 53.76% | 2487 / 46.24% | 1671 / 31.07% | 1331 / 24.75% | 2376 / 44.18% |

**Supplemental Figure 1.**
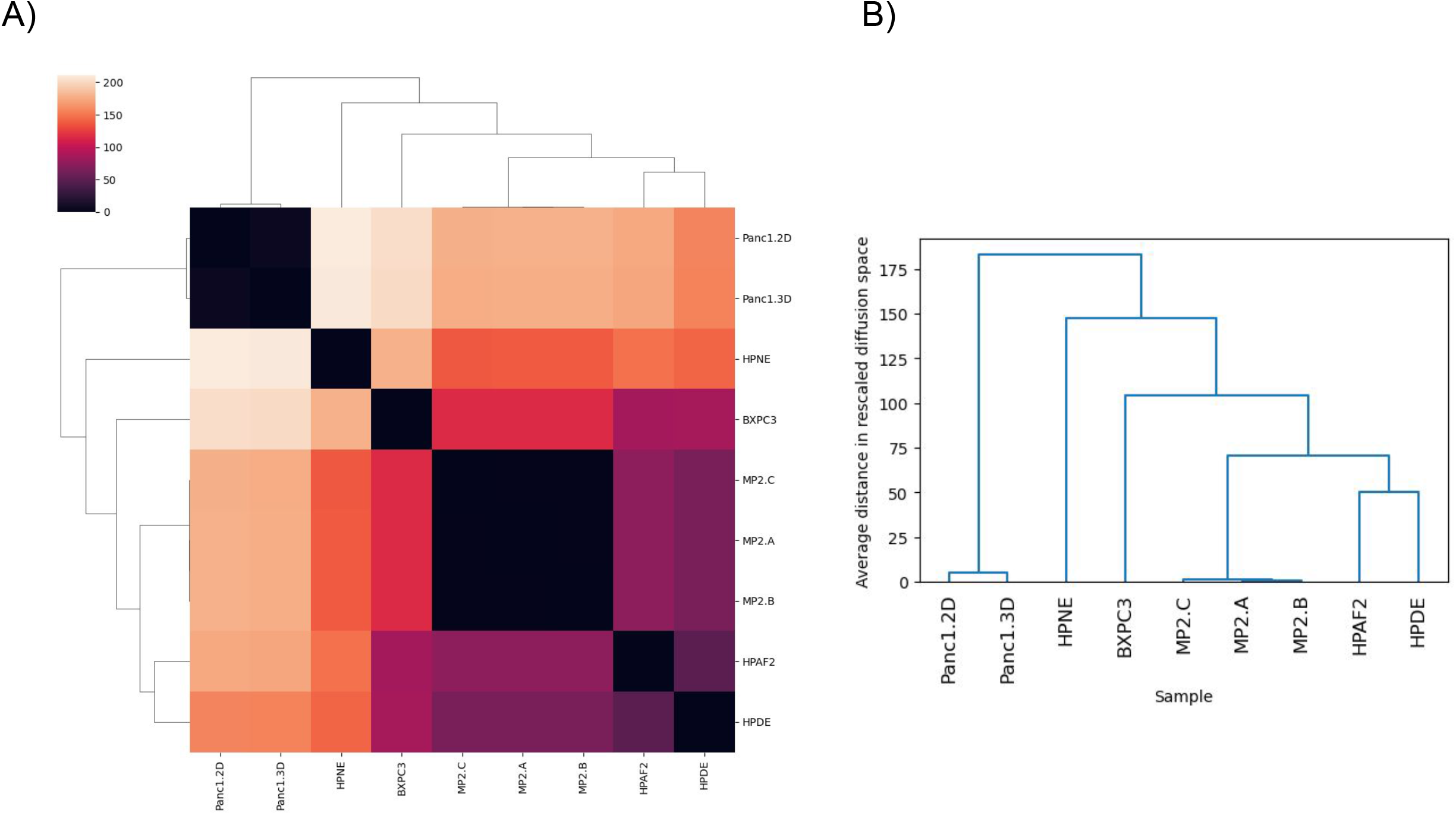
Phenotype mapping and cladogram analyses of cell lines depicting evolutionary relatedness between them. For the cladogram the x-axis indicates the sample names and y-axis indicates the phenotypic distances.

**Supplemental Figure 2.**
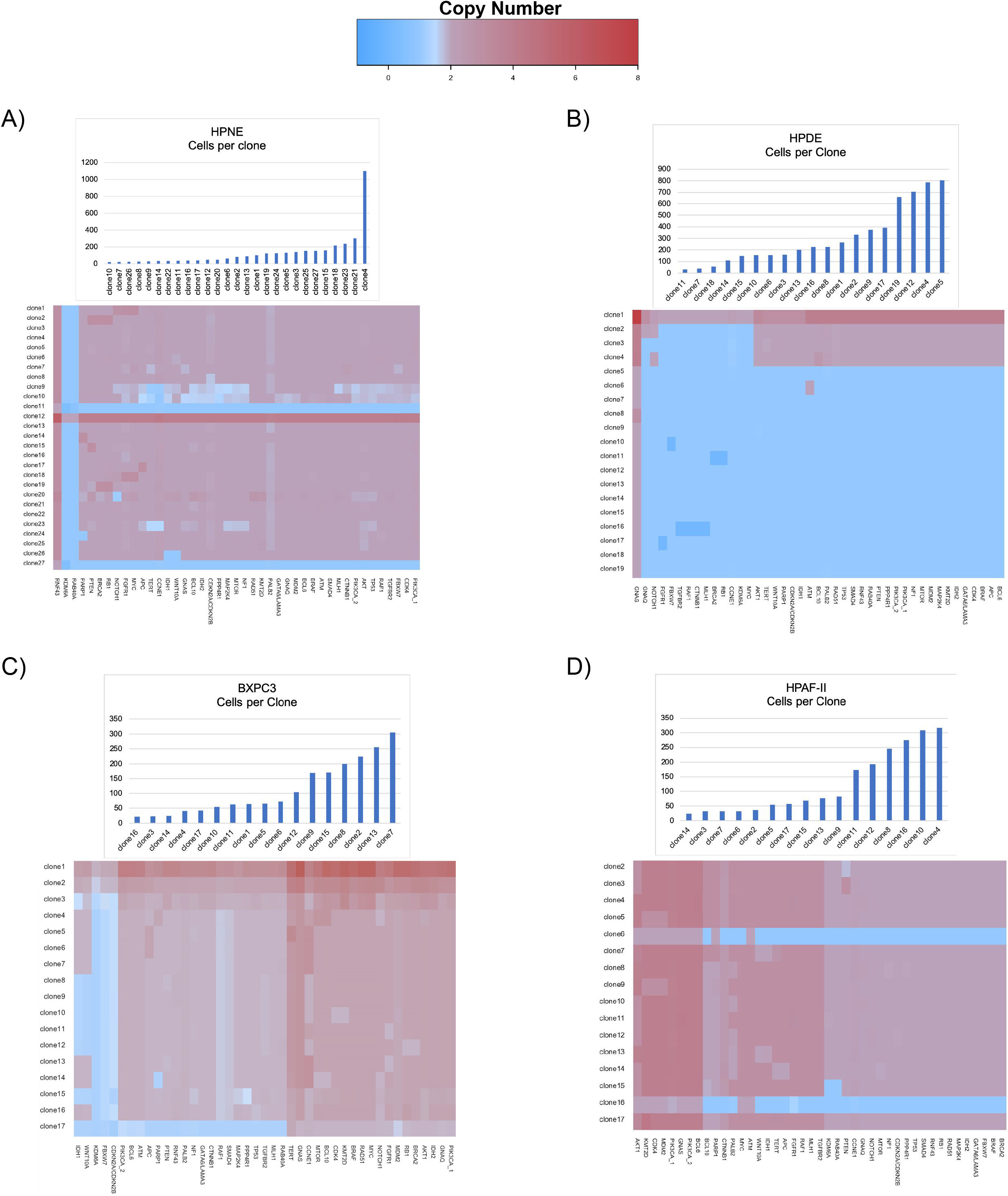
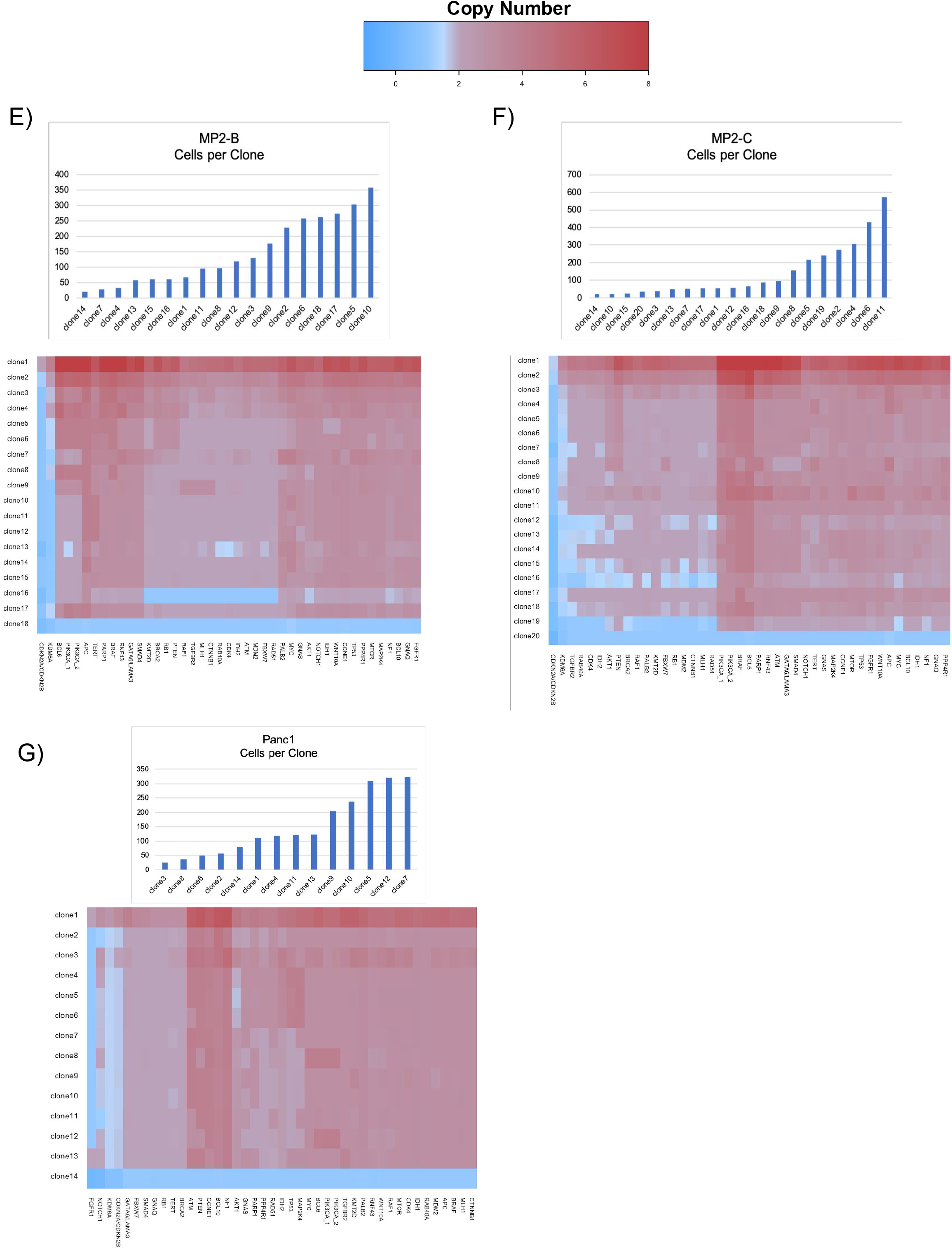
PDAC cell lines harbor clonal landscape with differential CNV events at oncogenic loci. For each cell line, histograms display the number of cells belonging to each genomically-unique clone. Heatmaps depict the number of copies per gene, per clone.

**Supplemental Figure 3.**
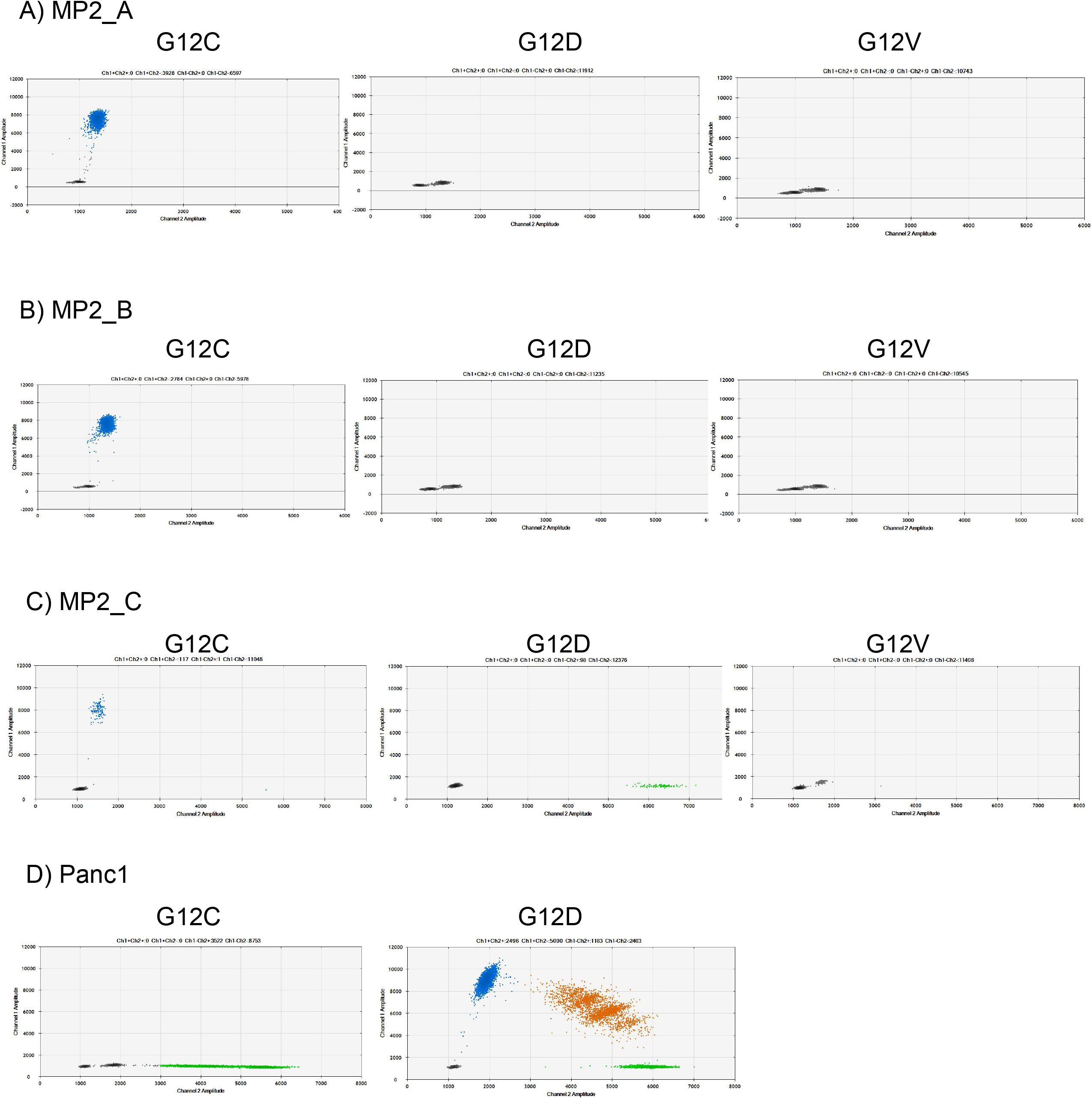
Hotspot KRAS mutations confirmed in PDAC cell lines. Droplet Digital PCR (ddPCR) results of KRAS probes for hotspot mutations G12C, G12D, and G12V applied to MP2 samples A-C and Panc1. **A-C)** Confirms same KRAS G12C mutation in all 3 cell lines. **D)** Panc1 ddPCR results indicating both KRAS G12D mutation.

**Supplemental Figure 4.**
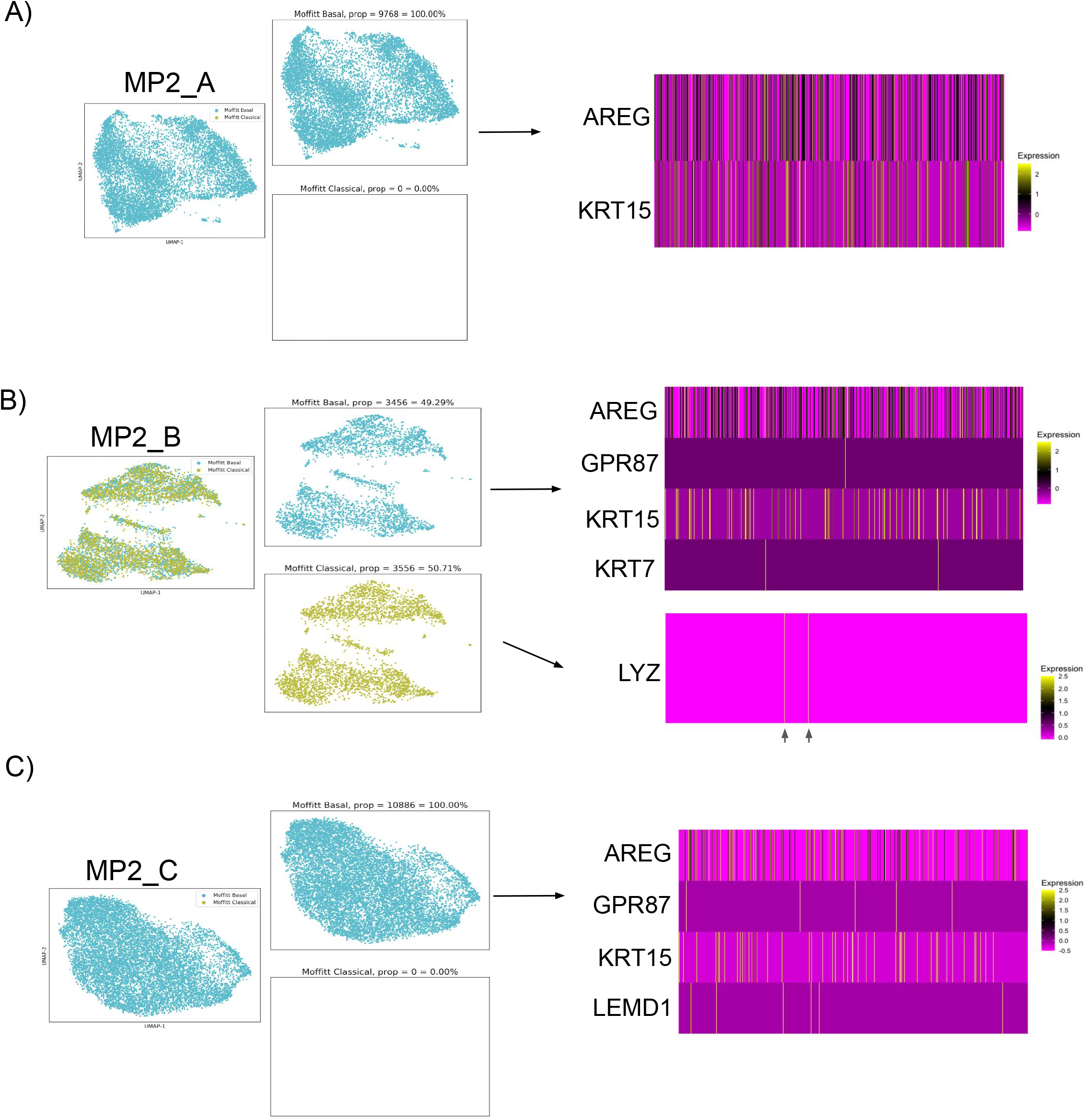
Expression of few genes dictates transcriptional subtyping of cell lines. Moffitt Classification, per gene, per cell across basal and classical subtypes in **A)** MP2-A, **B)** MP2-B, and **C)** MP-C displaying how a sparse expression of subtype genes across thousands of cells in each sample forces an artificial classification.

**Supplemental Figure 5:**
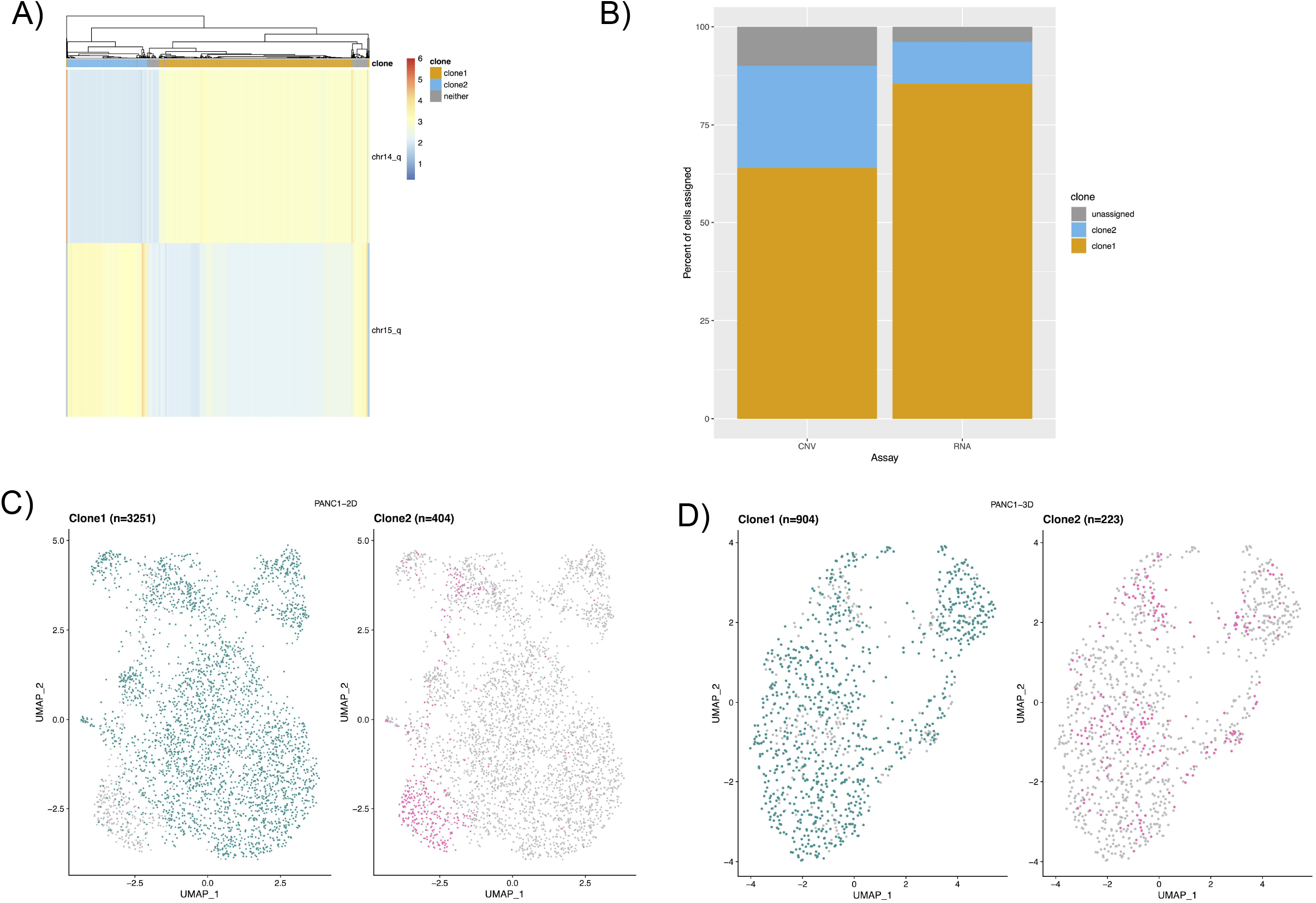
Genome-transcriptome mapping from parental Panc1 scCNVseq yields differential expression profiles between monolayer and spheroids. **A)** Clonealign results using the same genomic scCNV background. Clones demarcated by CNV events at chromosomes 14q and 15q. Cells per clonal group are along the x-axis, y-axis indicates CNV events used for subsetting clones. **B)** Clonealign scCNV clone correspondence to Panc1 2D scRNA data. **C)** UMAP of scCNV-derived clones mapped to corresponding scRNA-derived cells for Panc1 2D and **D)** Panc1 3D.

**Supplemental Figure 6.**
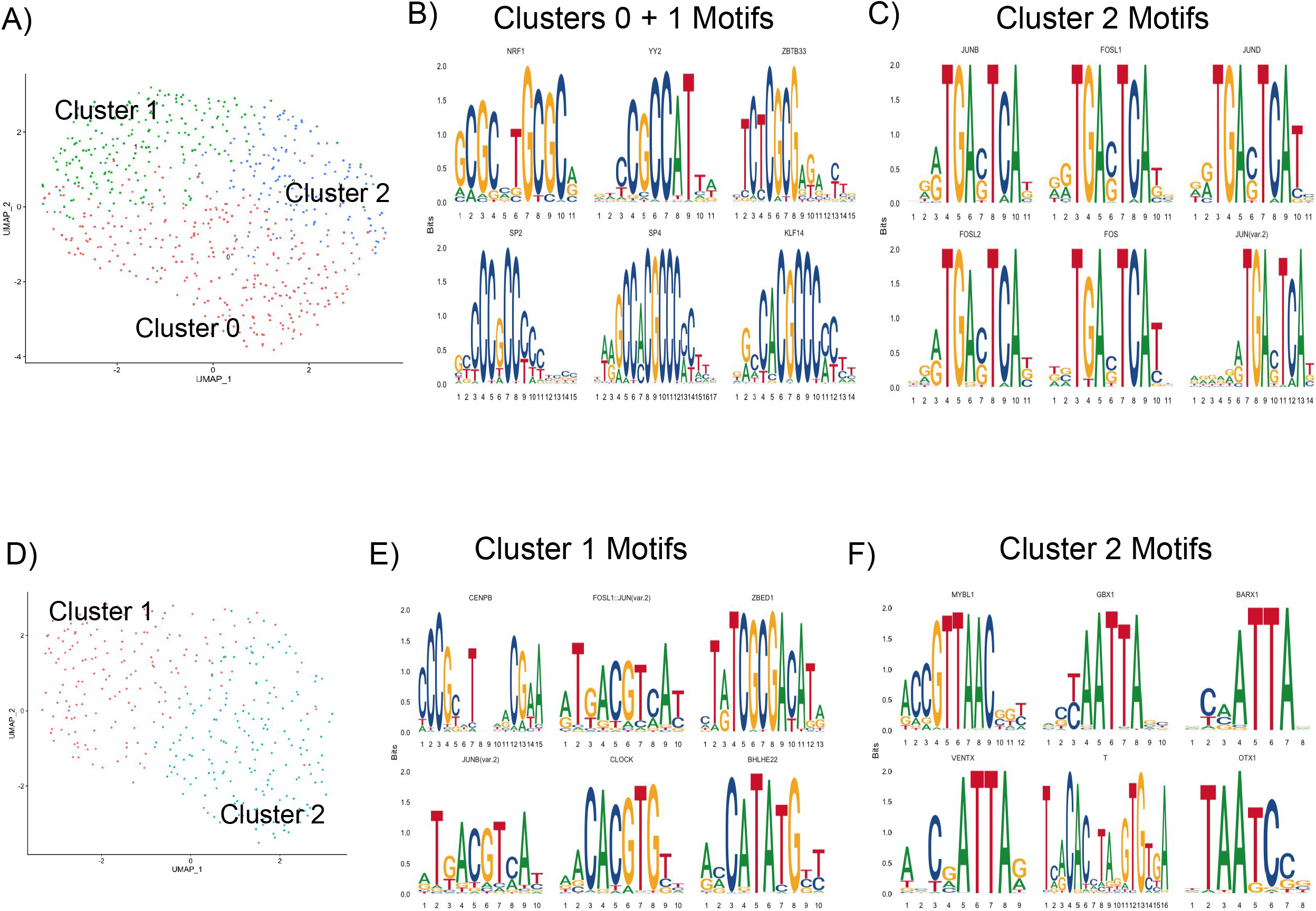
Chromatin modifications have transcriptional consequences in spheroid model. **A)** Individual analysis of Panc1 spheroid nuclei shows 3 predominating states of chromatin architecture (clusters 0-2) that correspond to unique enrichment patterns. **B)** Clusters 0 and 1 of Panc1 spheroid nuclei are enriched for SP motifs, NRF1, YY2, ZBTB33, and KLF14. **C)** Cluster 2 of Panc1 spheroid nuclei is enriched for FOS/Jun motifs. **D)** Individual analysis of Panc1 2D (monolayer) nuclei shows 2 predominating states of chromatin architecture (clusters 1, 2) that correspond to unique enrichment patterns. **E)** Cluster 1 of Panc1 monolayer nuclei is enriched for Fos/Jun motifs, ZBED1, CENPB, CLOCK, and BHLHE22. **F)** Cluster 2 of Panc1 monolayer nuclei is enriched for MYBL1, GBX1, BARX1, VENTX, T, and OTX1.

